# A comparison of convolutional neural networks and few-shot learning in classifying long-tailed distributed tropical bird songs

**DOI:** 10.1101/2023.07.25.550590

**Authors:** Ming Zhong, Jack LeBien, Marconi Campos-Cerqueira, T. Mitchell Aide, Rahul Dodhia, Juan Lavista Ferres

## Abstract

Biodiversity monitoring depends on reliable species identification, but it can often be difficult due to detectability or survey constraints, especially for rare and endangered species. Advances in bioacoustic monitoring and AI-assisted classification are improving our ability to carry out long-term studies, of a large proportion of the fauna, even in challenging environments, such as remote tropical rainforests. AI classifiers need training data, and this can be a challenge when working with tropical animal communities, which are characterized by high species richness but only a few common species and a long tail of rare species. Here we compare species identification results using two approaches: convolutional neural networks (CNN) and Siamese Neural Networks (SNN), a few-shot learning approach. The goal is to develop methodology that accurately identifies both common and rare species. To do this we collected more than 600 hours of audio recordings from Barro Colorado Island (BCI), Panama and we manually annotated calls from 101 bird species to create the training data set. More than 40% of the species had less than 100 annotated calls and some species had less than 10. The results showed that Siamese Networks outperformed the more widely used convolutional neural networks (CNN), especially when the number of annotated calls is low.

## I. Introduction

The 21st century is marked by the severe population decline of multiple taxonomic groups due to habitat loss, climate change, hunting, and introduced species changes (Sánchez-Bayo and Wyckhuys 2021; Rosenberg *et al*. 2019; Pacoureau *et al*. 2021; He *et al*. 2019; Spooner *et al*. 2018). To slow the loss of biodiversity, we urgently need to better understand how changes in climate and other environmental variables are affecting species distributions and abundances. Unfortunately, monitoring these state variables can be challenging, especially for species of greatest conservation concern, such as rare and endangered species.

Furthermore, in many ecosystems such as tropical rainforests, high species richness is made up of a relatively small number of common species and many rare species (Hubbell 2001). From a conservation or management perspective, these rare species are of utmost important, but up to now it has been a challenge to collect reliable long-term data for most of these species.

New tools (e.g., inexpensive audio recorders) and technologies (e.g., artificial intelligence) can greatly improve species identification and discovery. Most research on automating species identification in audio recordings has focused on producing algorithms specific to a single species (e.g., Aide *et al*. 2013). This approach limits the information that can be extracted from soundscapes, given that many species can be present in a single recording, particularly in species diverse habitats. In contrast, deep learning algorithms (e.g., neural networks) have been developed to identify multiple species (e.g., Zhong *et al*. 2020). These algorithms typically require a high number of annotated calls (i.e., training data) to achieve satisfactory accuracy. This is because the deep neural network models usually include millions of parameters and tend to overfit on small datasets, resulting in poor accuracy. To address the issue of limited training data, researchers have developed few-shot learning methods (Koch *et al*. 2015; Vinyals *et al*. 2016; Snell *et al*. 2017; Sung *et al*. 2018). These few-shot learning models take a contrastive learning approach using pairs or triplets of samples as training input. Since triplets of samples are compared in each training iteration - instead of comparing just one sample with its target label - the number of unique training samples effectively increases to the number of unique triplets in the training set (i.e., data augmentation).

Here we compare species identification results using two approaches: convolutional neural networks (CNN) and Siamese Neural Networks (SNN), a few-shot learning algorithm. The goal is to develop methodology that accurately identifies both common and rare species.

## II. Data

### A. Data Sources and Data Annotation

We collected more than 100,000 one-minute audio recordings from 99 sites on Barro Colorado Island (BCI), Panama in 2018 (Campos-Cerqueira *et al*. 2021).

These recordings were used to create a detection history of more than 100 species in the audio recordings through three steps. First, biological experts manually searched for species in recordings from 5:00 to 9:00 a.m. from each site and created a call template for each species. Second, in the RFCx-ARBIMON platform (Aide *et al*. 2013), we used the template matching algorithm by providing the system with the species-call template, a playlist of all recordings, and a correlation threshold (0.1). All detections above the correlation threshold were cropped and displayed for posterior validation. Third, the experts reviewed the template matching results and annotated the results as either positive or negative.

For the present study, we created a dataset of approximately 23,000 annotated calls from 101 bird species. The duration of most calls (87%) was less than 4 seconds, remaining 13% last between 4 and 7 seconds. The number of annotations varied greatly among species (Table 1). Eleven species had four or fewer annotations, while 55 species had more than 100.

**TABLE I:** Number of annotated calls per species.

| Number of Annotated Calls per species | Number of bird species |
| --- | --- |
| 1-4 | 11 |
| 5-99 | 35 |
| 100+ | 55 |

### B. Data for modeling

Using custom-written scripts in Python 3.7, Mel-spectrograms were produced from audio files (with NFFT = 1024 and 75% overlap, Hann window). Each mel-spectrogram was generated from a 4-s audio segment that contained either one or multiple annotated calls and was resized as 384 pixels by 384 pixels with RGB channels (i.e., colored Mel-spectrograms). During the annotation process, we only labelled a single species in each template-matching detection. As a result, for each extracted Mel-spectrogram, the presence or absence of only one species is labeled.

Given the long-tail distribution of the labeled data among all studied species, we grouped the 11 species with less than five annotations into one category; therefore, our model has 91 categories in total. The annotated data were randomly split into training, validation, and testing sets (which account for 64%, 16%, and 20% of the annotated data, respectively), and the model results were reported and evaluated on the testing set. To make a fair comparison, we used the same backbone architecture (DenseNet121) for both Convolutional Neural Networks (CNN) and Siamese Neural Networks (SNN).

## III. Methods

We assessed the performance of Convolutional Neural Networks (CNN) and a technique for few-shot learning, Siamese Neural Networks (SNN), to determine which best classified bird calls.

### A. Classification Models using Convolutional Neural Network (CNN)

Convolutional Neural Networks (CNN) have been widely used for image classification tasks, and their success has also been proven in bioacoustic classification applications (Bianco *et al*. 2019, Zhong *et al*. 2020, LeBien *et al*. 2020). Here we used the DenseNet architecture (Huang *et al.* 2016) as a baseline to classify the presence or absence of calls for each species in each 4-s spectrogram. DenseNet was explicitly developed to improve the negative effect on accuracy caused by the vanishing gradient in deep neural networks and has the advantage of improving feature propagation both in a forward and backward fashion. In a DenseNet architecture, the output feature-map of each layer is used as input for each subsequent layer, such that all layers are connected.

Since many deep neural network models have parameters in the order of millions, they heavily rely on big data to avoid overfitting (REF). However, almost 40% of species have less than 100 labeled calls in our annotated data. As an effective data-space solution to the problem of limited data, data augmentation refers to the techniques that attempt to artificially increase the size and quality of training datasets such that the models built using them may achieve higher accuracy.

Among various data augmentation methods for image processing, some basic ones include flips, rotations, shifts, noise injections, color space transformations, sharpening or blurring, and random erasing or cropping (REF). Specifically, for audio recordings, there are methods such as time-stretching, pitch shifting, and mixing multiple audio files (Salamon and Bello 2016). Beyond them, there are more advanced techniques, such as generative adversarial network (GAN)-based methods (Antoniou *et al.* 2017), which can generate synthetic images. For this model implementation, as our primary goal is to compare the performance between CNN and few-shot learning models, we did not apply advanced data augmentation techniques, but only two basic techniques instead to increase the size of data that can be used for model training: rotation (up to 5 degrees) and time-frequency shifting (width and height shifting up to 10% of the original spectrogram).

### B. Classification Models using Siamese Neural Network (SNN)

Siamese Neural Networks (SNN) (Koch *et al.* 2015) are a class of neural network architectures that contain two or more identical subnetworks. “Identical” here means having the same configuration with the same parameters and weights. Parameter updating is mirrored across both sub-networks. SNN focuses on learning image embeddings in the deeper layers that place the same classes close together. Hence, it can be used to measure the similarity of the inputs by comparing their feature vectors and deciding whether the two images belong to the same category or different categories.

Since training of Siamese networks involves pairwise learning, a cross-entropy loss cannot be used in this case. Instead, we used another loss function called triplet loss (Hoffer and Ailon, 2015). This is a loss function where an anchor (baseline) image is compared to a positive image (i.e., an image that is in the same category as the anchor image) and a negative image (i.e., an image that is in a different category as the anchor image). The distance from the anchor image to the positive image is minimized, and the distance from the anchor image to the negative image is maximized. As shown in formula (1), *D* (*x, y*) represents the distance between the learned vector representation of *x* and *y*. α is a margin term used to stretch the distance differences between similar and dissimilar pairs in the triplet. The remaining parameters represent the feature embeddings for the anchor (*a*), positive (*p*), and negative (*n*) images.

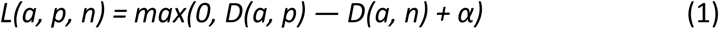

During the training process, an image triplet (anchor image, positive image, negative image) is fed into the model as a single sample (see Fig. 1). The distance between the anchor and positive images should be smaller than that between the anchor and negative images, indicating higher similarity between the anchor and positive images. An extensive training data set is needed for many deep learning models to achieve good performance. While this may not be practical in many real applications, the way how Siamese Networks make good use of all training examples to train embeddings enables these networks to learn from very little data.

**FIG. 1.**
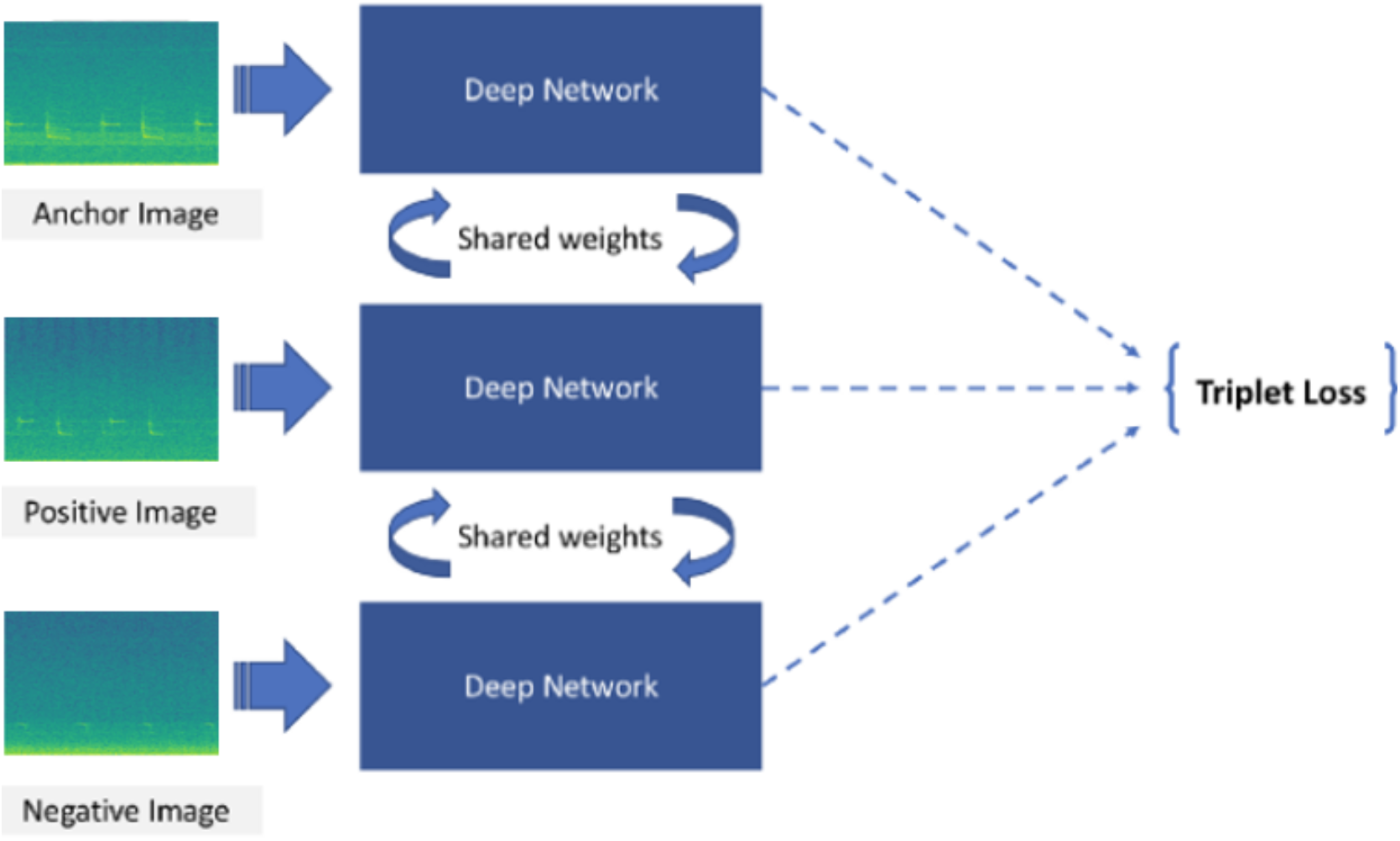
Architecture of Siamese Networks with triplet loss.

When triplets are generated for model training, as the training continues, some of the additional triplets are easy to deal with because their loss value is very small or even *0,* preventing the network from further improvement. A good training strategy would be to constantly “mine” out those difficult cases (i.e., triplets that distance between the anchor and positive image is larger than the distance between the anchor and negative image) in each epoch, based on the performance of the model’s current snapshot, so that the model will always have a certain percentage of challenging cases in the training loop from which it still struggles to tell the difference. This is similar to the triplet mining in FaceNet (Schroff *et al.* 2015).

After getting the embedding vector for each mel-spectrogram, we measured the similarity (i.e., L2 distance) for each mel-spectrogram in the test set with those in the training set and assigned the label to the closest species.

## III. Results

To evaluate the model performance on datasets with different categories and sizes, we reported the model performance when fitting on (1) all species and 2) rare species only. For each dataset, we reported the top-1, top-3, and top-5 accuracies (top-k accuracy is the accuracy where true class matches with any one of the k most probable classes predicted by the model), where the accuracies are calculated as an average of 5 independent runs.

CNN performs slightly better on top-1 accuracy for the overall dataset and common species for models fitted on the training data from the entire annotated data (Table 2). In comparison, SNN performs substantially better on three measures for rare species and has higher top-3 and top-5 accuracies for the overall dataset and common species.

**TABLE II:** Comparison of classification results for the CNN and SNN models. Results are shown for all species combined, common species (>=100 annotated calls), and rare species (<100 annotated calls). The highest performance for each measure and species subset is in bold type.

| Species | CNN |  |  | SNN |  |  |
| --- | --- | --- | --- | --- | --- | --- |
|  | Top-1 | Top-3 | Top-5 | Top-1 | Top-3 | Top-5 |
|  | Accuracy | Accuracy | Accuracy | Accuracy | Accuracy | Accuracy |
| All species | **89.48%** | 94.42% | 95.54% | 88.66% | **96.10%** | **97.35%** |
| common species | **91.22%** | 95.62% | 96.59% | 89.85% | **96.93%** | **98.08%** |
| rare species | 67.12% | 79.02% | 81.90% | **73.37%** | **85.46%** | **87.91%** |

When the analyses are restricted to the species with small training sets (<100 annotations), the difference in the performance of the two models is even more dramatic. The accuracy of CNN decreases to a much larger extent than that of SNN (Table III).

**TABLE III:** Model results for classifying the presence of rare species (i.e., fewer than 100 annotations per species) for the CNN and SNN models. The highest performance for each measure and species subset is in bold type.

| Species | CNN |  |  | SNN |  |  |
| --- | --- | --- | --- | --- | --- | --- |
|  | Top-1 | Top-3 | Top-5 | Top-1 | Top-3 | Top-5 |
|  | Accuracy | Accuracy | Accuracy | Accuracy | Accuracy | Accuracy |
| 5 – 100 annotated calls<br>(Includes 35 species) | 83.25% | 90.25% | 92.21% | **85.77 %** | **93.19%** | **95.40%** |
| 5 – 20 annotated calls<br>(Includes 13 species) | 53.69% | 67.89% | 75.79% | **73.16%** | **90.00%** | **93.69%** |
| 5 – 10 annotated calls<br>(Includes 7 species) | 35.29% | 64.71% | 82.35% | **60.00%** | **80.00%** | **90.59%** |

## IV. Discussion

We built models to classify common calls of 101 bird species in the Barro Colorado Island (BCI), Panama. In comparison to CNNs, which have been successfully used to classify multiple species in field audio recordings, SNN achieved better performance in this study, when the number of training samples is limited.

The original manually annotated data includes detections, either positive or negative, indicating the corresponding species’ presence or absence. However, we only used the positively annotated detections in the modeling process due to computational constraints. On the other hand, the negatively labeled detections may be used as “difficult cases” when constructing triplets while training Siamese Networks. With the model that has been trained with only positive detections, we scored all the positive and negative detections in the test set with the hope that the model was able to distinguish cases of presence and absence for each species. The positive samples should have smaller distances (i.e., higher similarities) to the same species in the training set for each species. We can also find the globally optimal distance threshold that can distinguish between positive and negative detections. Our results show that for this dataset, the globally optimal threshold when detecting a species’ call is around 0.3 (Figure 2). The smaller the distance, the higher confidence we have that the classified species’ call is correct. Further, as different species’ calls have different similarities or uniqueness, it may be even better to choose species-wise distance thresholds.

**FIG. 2.**
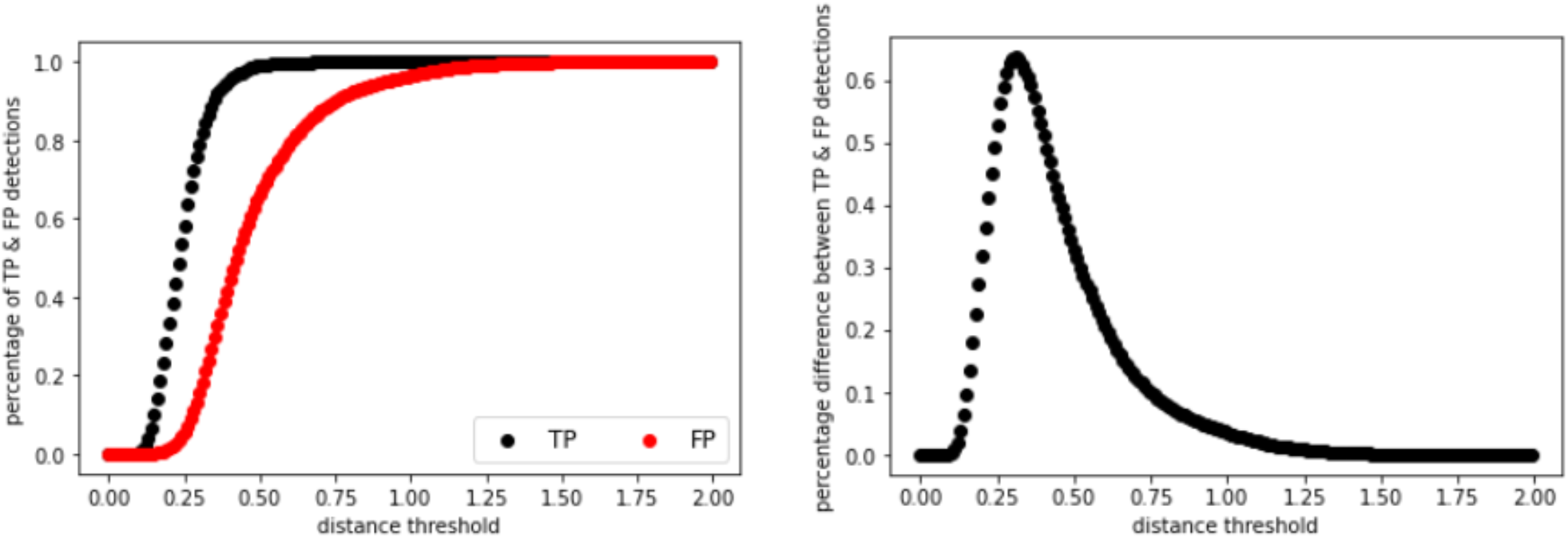
Comparison between the positive (TP) and negative (FP) detections at different distance thresholds.

